# Endosymbiotic *Rickettsiella* causes cytoplasmic incompatibility in a spider host

**DOI:** 10.1101/2020.03.03.975581

**Authors:** Laura C. Rosenwald, Michael I. Sitvarin, Jennifer A. White

## Abstract

Many arthropod hosts are infected with bacterial endosymbionts that manipulate host reproduction, but few bacterial taxa have been shown to cause such manipulations. Here we show that a bacterial strain in the genus *Rickettsiella* causes cytoplasmic incompatibility (CI) between infected and uninfected hosts. We first surveyed the bacterial community of the agricultural spider *Mermessus fradeorum* (Linyphiidae) using high throughput sequencing and found that individual spiders can be infected with up to five different strains of maternally-inherited symbiont from the genera *Wolbachia, Rickettsia*, and *Rickettsiella*. The *Rickettsiella* strain was pervasive, found in all 23 tested spider matrilines. We used antibiotic curing to generate uninfected matrilines that we reciprocally crossed with individuals infected only with *Rickettsiella*. We found that only 13% of eggs hatched when uninfected females were mated with *Rickettsiella*-infected males; in contrast, at least 83% of eggs hatched in the other cross types. This is the first documentation of *Rickettsiella*, or any Gammaproteobacteria, causing CI. We speculate that induction of CI may be much more widespread among maternally-inherited bacteria than previously appreciated. Further, our results reinforce the importance of thoroughly characterizing and assessing the inherited microbiome before attributing observed host phenotypes to well-characterized symbionts such as *Wolbachia*.

## Introduction

Maternally transmitted endosymbiotic bacteria are common in arthropods, and are estimated to infect most arthropod species [1–4]. These inherited endosymbionts frequently manipulate host reproduction to promote symbiont transmission and spread in the host population [5]. The most prevalent and well-studied symbiont-driven manipulation is cytoplasmic incompatibility [5–8]. Cytoplasmic incompatibility (CI) is a conditional sterility phenotype in which crosses between infected males and uninfected females result in offspring mortality, sabotaging the production of uninfected progeny. This phenotype functionally increases the proportion of infected females in the host population over time [5]. Cytoplasmic incompatibility influences host population dynamics and gene flow, acting both as a gene drive mechanism that spreads traits through a population [9] and as a reproductive barrier that can enforce reproductive isolation and contribute toward speciation [10,11]. In application, CI has been used to locally repress pest populations as a form of sterile insect technique [12], and also as a gene drive/defensive mechanism to reduce disease transmission by mosquitoes [13,14].

To date, few bacterial taxa have been shown to cause CI. The most widespread and well-investigated are members of the genus *Wolbachia* (Phylum Proteobacteria, Class Alphaproteobacteria, Order Rickettsiales), which infects many arthropod species [1,8,15,16]. Recently, another unnamed strain in the Rickettsiales has also been suggested to induce CI [17]. Finally, members of the more distantly related *Cardinium* genus (Phylum Bacteroidetes) also induce CI, although using a mechanism that appears to be independently evolved from *Wolbachia* [18]. No other bacterial taxa have been documented to cause CI to date.

In the present study, we provide evidence that CI can be caused by a strain of *Rickettsiella* (Phylum Proteobacteria, Class Gammaproteobacteria, Order Legionellales), a bacterium that is not closely related to either *Wolbachia* or *Cardinium* [19]. We began by characterizing the microbiome of *Mermessus fradeorum* (Araneae: Linyphiidae), an agricultural spider that had been previously shown to be infected by multiple *Wolbachia* and *Rickettsia* (Phylum Proteobacteria, Class Alphaproteobacteria, Order Rickettsiales) symbionts, and to exhibit CI [20]. Upon discovery of an unexpected *Rickettsiella* symbiont that was pervasive throughout our sampled population, we proceeded to experimentally document that *Rickettsiella* causes CI in *M. fradeorum*. This is the first demonstration of CI by a Gammaproteobacteria, which suggests that the taxonomic distribution of bacteria that cause CI may be wider than previously appreciated, and could have important implications for the population genetics, biodiversity, and evolution of arthropod hosts.

## Methods

### Study System

We initially collected *Mermessus fradeorum* from alfalfa at the University of Kentucky’s Spindletop Research Farm (38.127, −84.508). We placed individual spiders into deli cups (6cm diam.) with hydrated plaster in the bottom for humidity control, and maintained them in the laboratory at 22°C 24hr dark conditions. Spiders collected as juveniles were fed symbiont-free collembola (*Sinella curviseta*) until maturity. Adult spiders were fed fruit flies (*Drosophila melanogaster*) at 2-3 day intervals; these flies were infected with a distinct strain of *Wolbachia* (MLST ST1) [21] that was never detected in our microbiome survey or diagnostic assays. We specifically tested whether infected prey could result in false-positive infection diagnoses in the spiders, and found that even an hour after feeding on *Wolbachia*-infected flies, uninfected *M. fradeorum* did not test positive for *Wolbachia* (Electronic supplemental material figure S1). We initiated matrilines from individual field-collected gravid females, or by mating field-collected females to males within the laboratory.

### Microbiome Survey

To evaluate endosymbiont diversity in the population of *M. fradeorum*, we characterized the microbiome of 23 matrilines that were initiated between August and October 2016. For each matriline, we evaluated the field-collected female specimen when possible; if the field-collected female died of natural causes (n = 8), we instead randomly selected an adult female F1 offspring. We withheld food from each spider for at least five days before preserving the specimen in 95% ethanol. We surface-sterilized each specimen with a series of washes with 0.5% bleach and PCR water. We then extracted total DNA from each individual specimen using DNEasy Blood and Tissue kits (Qiagen, Germantown, MD) according the manufacturer’s instructions. Using PCR with dual-indexed 515F/806R primers [22], we amplified the V4 region of bacterial 16S from each specimen and visualized an aliquot of the resulting product on a 1% agarose gel stained with GelRed (Biotium). After verifying the presence of a strong band of the expected product size, we included a 1µl aliquot of the corresponding product in a multiplexed library for sequencing. This library also included samples from aphids and parasitoid wasps that served as controls (see below). After purification using GenCatch PCR Cleanup kit (Epoch Life Sciences, Missouri City, TX) we sequenced the library on an Illumina Miseq 2500 instrument at the University of Kentucky’s core sequencing facility.

Sequences were demultiplexed, trimmed and quality filtered within BaseSpace (Illumina) then imported into qiime2 (v2017.11) [23] using a manifest. We conducted additional quality control using deblur [24], implemented in qiime2 using default parameters and a trim length of 251 bases. To determine the taxonomic placement of each operational taxonomic unit (OTU), we used a naïve Bayes classifier trained on the V4 region of the Greengenes 13_8 99% OTUs reference database [25].

The most prevalent OTU in the library, a strain of *Rickettsiella*, constituted 46% of total reads and was associated with every component specimen in the multiplexed sample, albeit at low prevalence for non-*M. fradeorum* samples. We inferred that index swapping may have occured between samples that shared either forward or reverse indices, resulting in reads that were misallocated to the wrong sample [26]. To validate that *Rickettsiella* was genuinely present in all *M. fradeorum* samples, we used *Rickettsiella*-specific primers to diagnostically test for the presence of *Rickettsiella* in the 23 matriline samples, as well as the 8 controls (4 *Aphis craccivora* (aphid) specimens and 4 *Habrobracon hebetor* (parasitoid wasp) specimens). Diagnostic primers and protocols followed Duron et al. [19]; samples that tested negative for *Rickettsiella* were rescreened at least two more times, and additionally tested for extraction quality via diagnostic PCR of a segment of the arthropod COI gene [4].

To assess vertical transmission of *Rickettsiella*, we included specimens from several generations in the microbiome dataset. In total, 148 mother-offspring pairs from the 23 matrilines (representing 83 *Rickettsiella*-infected mothers) were initially included in the dataset. As a quality control measure for the final dataset, we retained only pairs in which both mother and offspring yielded more than 5000 reads (117 pairs, representing 73 different mothers, sometimes with multiple offspring). To be scored positive, the *Rickettsiella* read number for a given microbiome needed to exceed the upper 95% confidence boundary of *Rickettsiella* reads found in the eight control samples (145 reads).

### Rickettsiella *Cytoplasmic Incompatibility Assay*

To assess the ability of *Rickettsiella* to cause CI, we first created a population of uninfected spiders using antibiotics. Spiderlings originating from seven infected matrilines were treated with a combination of tetracycline (0.1%) and ampicillin (0.1%) by a fine mist spray daily until sub-adulthood [20,27]. Upon maturity, treated spiders were mated to one another, and a subset of those offspring were diagnostically tested for endosymbionts (*Wolbachia, Rickettsia, Rickettsiella)* [4,20]. Sibships that appeared negative for all endosymbionts were used to initiate the putatively uninfected population for experimental assays, with the understanding that not all spiders within this group would be truly cleared of infection. Antibiotic treatment often destabilizes symbiont transmission to offspring, resulting in inconsistent infection status among sibships and/or low titer infections that rebound in subsequent generations [28,29]. It is common practice to treat hosts with antibiotics for several generations to ensure symbiont clearance, but due to the labor intensive rearing of cannibalistic spiders, here we chose instead to diagnostically test all parental spiders to properly classify mating cross types, as described below. Spiders used in experiments were 1, 2, or 3 generations removed from antibiotic treatment.

To determine if *Rickettsiella* was causing CI in *M. fradeorum*, we reciprocally mated uninfected spiders with spiders that were infected with only *Rickettsiella*. We randomly assigned virgin spiders to mate with a partner that we expected to be either *Rickettsiella*-infected or uninfected, ensuring that siblings and cousins were not paired with one another. Males and females were paired for 8 hours, and were observed at approximately 15 minute intervals for the first two hours and periodically thereafter. If the pair was not observed to mate, they were excluded from the trial. Females were then allowed up to six weeks to lay three egg masses; females that failed to lay three egg masses within this time period (n=2) were excluded from the sample set. Egg masses were checked daily for spiderlings. Once spiderlings were observed, we allowed 24 hours for emergence, then dissected the egg mass to quantify unhatched eggs. Egg masses hatched within a mean ± SE of 14.1 ± 0.3 days. If no spiderlings were observed within 20 days, the entire egg mass was scored as unhatched, and dissected to count the number of unhatched eggs. Throughout the experiment, we found a small number of spiderlings (n=30) that hatched but died prior to exiting the egg mass. These unemerged spiderlings were excluded from the data set, as they accounted for only 2% of all eggs.

The infection status of all parents in the experiment was validated using diagnostic PCR for *Wolbachia, Rickettsia*, and *Rickettsiella* [4,19,20]. Final classification of all matings was based on these diagnostic results, rather than initial expectations based on sibling or parental infection status. Matings were reclassified if *Rickettsiella* presence/absence in either parent deviated from expectation, and were excluded altogether if either parent tested positive for a symbiont other than *Rickettsiella* (Electronic supplementary material table S1). The final distribution of mating cross types across four trial dates included 18 pairings in which both female and male were infected with only *Rickettsiella* (+/+), 14 in which the female was *Rickettsiella*-infected and the male was uninfected (+/-), 12 in which the female was uninfected and the male was *Rickettsiella*-infected (-/+), and 8 in which both female and male were uninfected (-/-).

We compared total eggs laid among treatments using ANOVA (JMP) after verifying that model assumptions of normality and homoscedacity were not violated. We compared egg hatch among treatments using logistic regression (Arc v. 1.06) with Williams correction [30] to account for moderate overdispersion. To verify that generations removed from antibiotic treatment (1, 2, or 3 generations) did not influence experimental outcome, we considered a separate model that incorporated this factor, but found no significant effect (Wald = −0.93, p = 0.35). Generation from antibiotic treatment was therefore not included in the final model. We additionally tested whether our diagnostic reclassification and exclusion criteria affected our conclusions by running the statistical tests on alternative datasets in which we 1) completely ignored our diagnostic results and had no reclassification, 2) reclassified according to *Rickettsiella* infection but ignored the presence of other symbionts (no exclusions), or 3) excluded all matings that did not fit with their initial expectations (exclusions instead of reclassifications). All of these variant analyses yielded statistically significant results that concur with the analysis of the original dataset presented below.

## Results

### Microbiome Survey

The most notable feature of the *M. fradeorum* microbiome was the pervasiveness of a single strain of *Rickettsiella*, present in all 23 matrilines (figure 1). Mean ± SE reads per sample was 22,054 ± 2,739, of which 16,172 ± 2,109 (80 ± 6%) were *Rickettsiella*. In most specimens (20/23), *Rickettsiella* constituted the majority of reads, often representing >95% of reads (figure 1, Electronic supplementary material tables S2-S3). Proportional representation of *Rickettsiella* was lower in samples that were also infected with *Rickettsia* (7 samples) and/or up to 3 strains of *Wolbachia* (8 samples), but still constituted half the reads returned for this subset of samples (50 ± 9%). In contrast, aphid and parasitoid wasp control samples that were included in the same sequencing run had few reads of this *Rickettsiella* strain (115 ± 15 reads/sample, constituting 0.3 ± 0.05% of reads/sample). Diagnostic validation confirmed the presence of *Rickettsiella* from all 23 matriline samples, and the absence of *Rickettsiella* from the aphids and parasitoid wasps. We observed 100% vertical transmission efficiency in the unmanipulated mother-offspring pairs in the dataset (Electronic supplementary material table S4). Across 117 pairs, all offspring of *Rickettsiella*-infected mothers were themselves infected with *Rickettsiella*.

**Figure 1.**
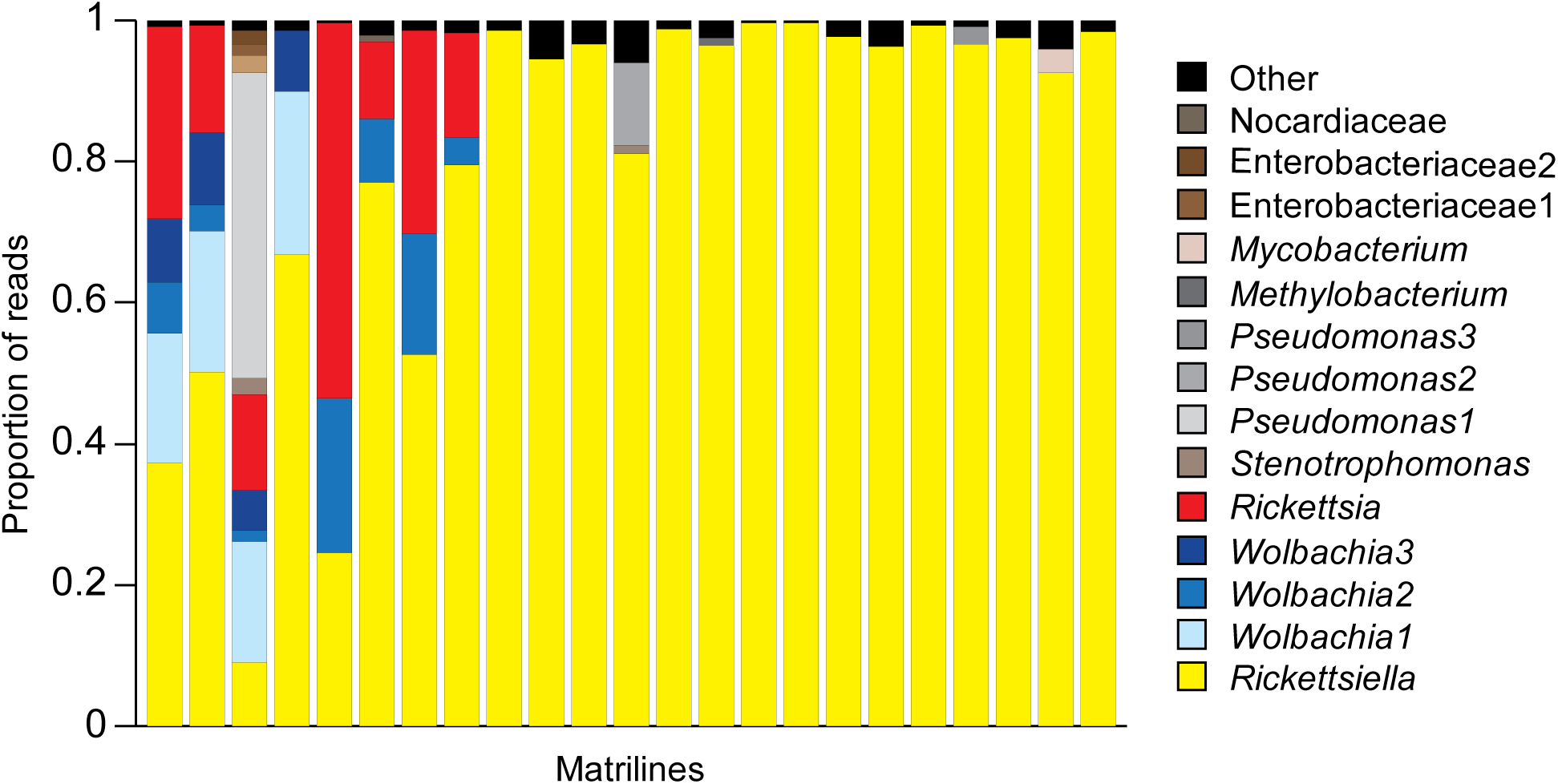
Proportional composition of 16S bacterial reads from 23 adult female *Mermessus fradeorum* spiders. All spiders were either field collected or adult offspring of different field collected mothers. Each shade represents a different bacterial strain; different strains from the same genus are designated by different numbers in the legend. Bacteria known to be maternally transmitted are brightly colored (yellow, blue, red), other strains are shades of gray and brown. Strain sequences, read numbers, and composition of the “Other” category available in Electronic supplemental material Tables S2 and S3.

### Reproductive Manipulation Assay: Cytoplasmic Incompatibility

When we reciprocally mated spiders bearing only *Rickettsiella* with those from matrilines that had been cured of all facultative symbionts, we found a large difference in the number of hatchlings produced. Total egg number was consistent and not significantly different across treatments (figure 2, Electronic supplementary material table S1; ANOVA: F3,48 = 0.56, p = 0.644), but the odds that these eggs hatched was significantly lower for uninfected females mated to *Rickettsiella*-infected males, relative to all three other treatments, which did not differ significantly from one another (χ2 = 119, d.f. = 3, p < 0.001). Only 13.2 ± 3.8% of eggs hatched in the incompatible cross between uninfected females and *Rickettsiella*-infected males, compared to 99.7 ± 0.2% of eggs when both parents were infected, 91.3 ± 4.6% from *Rickettsiella*-infected females mated with uninfected males, and 83.9 ± 12.2% when both parents were uninfected.

**Figure 2.**
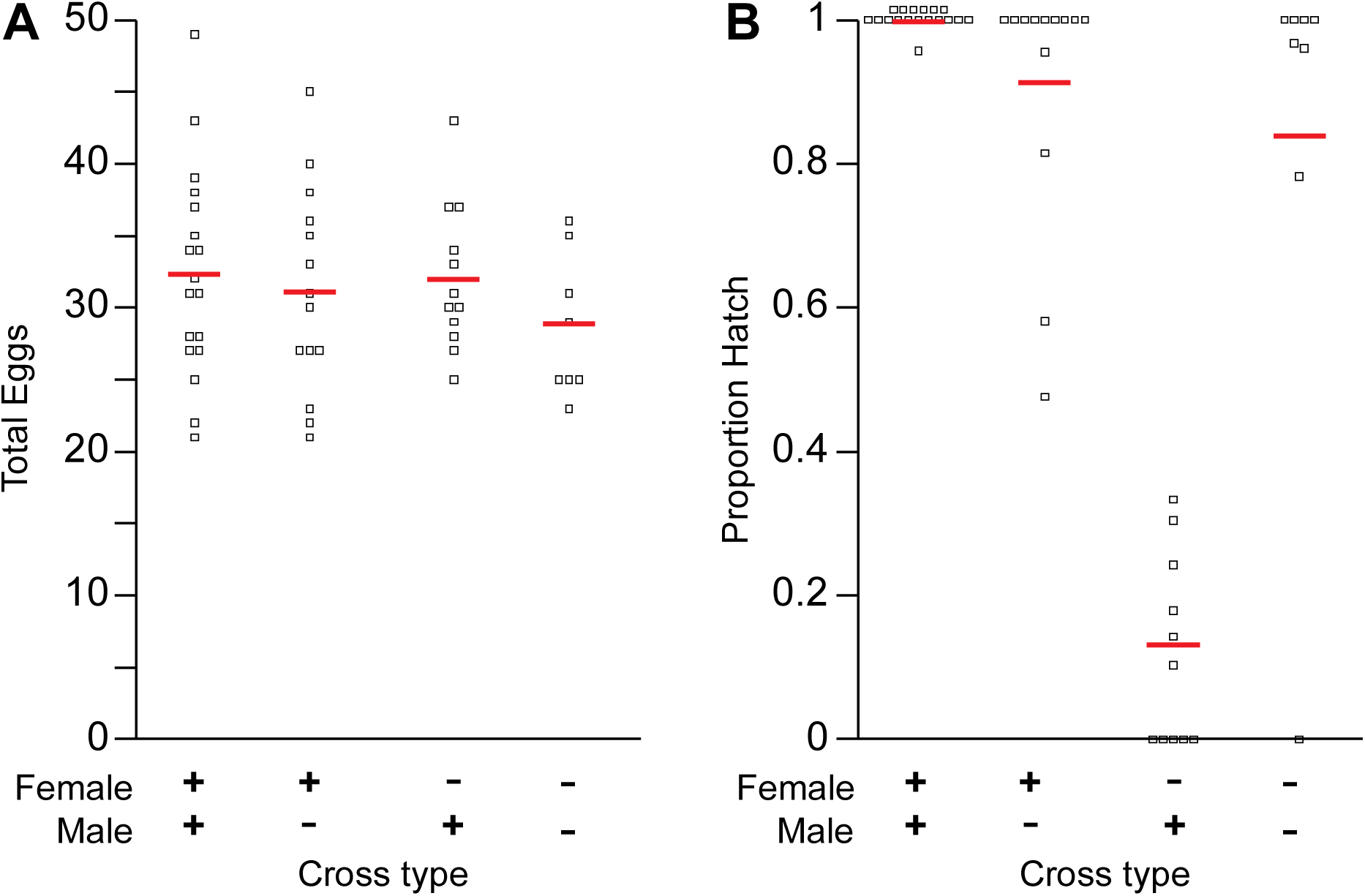
Total eggs laid (A) and proportion eggs that hatched (B) in reciprocal crosses between *Mermessus fradeorum* females and males that varied in *Rickettsiella* infection status. Each point represents the outcome from a single cross, red horizontal bars represent mean value per treatment. Points above 1.0 in panel B are 1.0.

## Discussion

We found that the spider *Mermessus fradeorum* is infected by up to five strains of bacterial symbiont, including a strain of *Rickettsiella* that was present in 100% of our field-collected matrilines. We then generated uninfected matrilines, and used crossing experiments to document that this *Rickettsiella* strain induced cytoplasmic incompatibility in the spider host. Previous examples of CI-causing bacteria have been restricted to *Wolbachia* and closely-allied bacteria in the Alphaproteobacteria, and *Cardinium* in the Bacteroidetes [6,7,17,31]. Ours is the first demonstration of a Gammaproteobacteria, *Rickettsiella*, causing CI.

Previous studies with this spider species diagnostically documented three endosymbiotic bacterial strains (two *Wolbachia* and one *Rickettsia*) associated with bacterially-induced reproductive manipulations, but could not attribute causality to individual bacterial strains [20]. The present microbiome study confirmed the presence of these same three symbiont strains at similar field frequencies as the previous work, and additionally revealed a third strain of *Wolbachia* along with the ubiquitous *Rickettsiella*. We were able to screen archived specimens from the previous study [20] and found that the CI-inducing matriline from that study was in fact co-infected with *Rickettsiella* and one strain of *Wolbachia* (L. Rosenwald, unpublished results). It is therefore likely that the CI observed in this previous study was at least partially attributable to *Rickettsiella*, although both co-infecting symbionts may have contributed to the CI phenotype [32]. Focus on taxa that are well-established as reproductive manipulators, such as *Wolbachia*, may lead researchers to overlook co-infecting bacteria that cause or contribute to manipulated host reproductive phenotypes, as co-infections are common in arthropods [1,3].

The mechanism by which *Rickettsiella* induces CI is unknown. In *Wolbachia*, CI is induced by phage-encoded operons in which one gene modifies the sperm of infected males and the other rescues the modification in the cytoplasm of the egg of infected females [33,34]. In *Cardinium*, the mechanism to induce CI appears to be independently evolved from *Wolbachia*, and is not phage associated [18]. No previously described *Rickettsiella* strains have been documented to cause CI, but CI-induction has been hypothesized based on genetic analysis of a *Rickettsiella* strain in fleas [35]. Strains of *Rickettsiella* infect a wide range of arthropods [19,36], sometimes as a pathogen [36], sometimes a mutualist [37], and sometimes as a potential reproductive manipulator [38]. Speculatively, it is possible that horizontal gene transfer among co-infecting symbionts may be responsible for *Rickettsiella’s* ability to induce CI [39,40], particularly given the high frequency of symbiotic co-infection in spiders in general [1,3,4,41] and *M. fradeorum* in particular.

Our data also reinforce the perception that spiders can harbor exceptionally diverse communities of co-infecting symbionts, rivaling the most symbiont-rich inherited communities among insects [4,42,43]. The discovery of an unexpected reproductive manipulator within such a community highlights the hazard of focusing only on well-characterized manipulative taxa (e.g. *Wolbachia*) when reproductive anomalies are observed in a host population. Our data emphasize the need for careful characterization of the entire inherited microbiome before attributing reproductive phenotypes to particular bacterial strains.

Reproductive manipulation by endosymbionts, through mechanisms such as CI, can influence arthropod behavioral ecology [44], population dynamics [45], and evolutionary trajectory [46]. As the known roster of symbionts capable of undertaking these manipulations increases, the potential for understanding and potentially exploiting their effects on the biology of their hosts increases as well. CI is currently being used as part of a symbiont-based strategy to decrease disease transmission in mosquitoes [12]; newly discovered CI-inducing symbionts, such as described in the present work, have similar potential in a wider range of arthropod hosts. In particular, *Wolbachia* is rare or absent in ticks (Class Arachnida, Order Acari), but strains of *Rickettsiella* are routine [19,38,47,48]. It would be worthwhile to evaluate the function of *Rickettsiella* and other members of tick microbiomes, similarly to what we have done with *Rickettsiella* in the spider *M. fradeorum*, as the potential opportunities for symbiont-based pest control and vector management are substantial.

## Data accessibility

The datasets supporting this article have been uploaded as part of the electronic supplementary material (Electronic supplementary material figure S1, tables S1-S4). High throughput sequencing data is available in the NCBI Sequence Read Archive under accession number PRJNA604923.

## Supporting information

Electronic supplemental material figure S1

Electronic supplementary material table

## Acknowledgements

We thank M. Curry and A. Styer for their previous work on this system, K. Welch for his *M. fradeorum* collections, and K. Athey for confirmation and initiation of *M. fradeorum* matrilines. We additionally thank S. Ferrell, A. Lindsey, K. O’Hearn and G. Moses for laboratory assistance and two anonymous reviewers for feedback on a previous version of this paper.

## References

1. Duron O, Bouchon D, Boutin S, Bellamy L, Zhou L, Engelstädter J, Hurst GD. 2008 The diversity of reproductive parasites among arthropods: *Wolbachia* do not walk alone. BMC Biol. 6, 1–12. (doi:10.1186/1741-7007-6-27)

2. Weinert LA, Araujo-Jnr EV., Ahmed MZ, Welch JJ. 2015 The incidence of bacterial endosymbionts in terrestrial arthropods. Proc. R. Soc. B Biol. Sci. 282, 3–8. (doi:10.1098/rspb.2015.0249)

3. Zhang L, Yun Y. 2018 Insights into the bacterial symbiont diversity in spiders. Ecol. Evol. 8, 4899–4906. (doi:10.1002/ece3.4051)

4. White JA, Styer A, Rosenwald LC, Curry MM, Welch KD, Athey KJ, Chapman EG. 2019 Endosymbiotic bacteria are prevalent and diverse in agricultural spiders. Microb. Ecol. 79, 472–481. (doi:10.1007/s00248-019-01411-w)

5. Engelstädter J, Hurst GDD. 2009 The ecology and evolution of microbes that manipulate host reproduction. Annu. Rev. Ecol. Evol. Syst. 40, 127–149. (doi:10.1146/annurev.ecolsys.110308.120206)

6. O’Neill SL, Giordano R, Colbert AM, Karr TL, Robertson HM. 1992 16S rRNA phylogenetic analysis of the bacterial endosymbionts associated with cytoplasmic incompatibility in insects. Proc. Natl. Acad. Sci. 89, 2699–2702. (doi:10.1073/pnas.89.7.2699)

7. Hunter MS, Perlman SJ, Kelly SE. 2003 A bacterial symbiont in the Bacteroidetes induces cytoplasmic incompatibility in the parasitoid wasp *Encarsia pergandiella*. Proc. R. Soc. B Biol. Sci. 270, 2185–2190. (doi:10.1098/rspb.2003.2475)

8. Werren JH, Baldo L, Clark ME. 2008 *Wolbachia*: Master manipulators of invertebrate biology. Nat. Rev. Microbiol. 6, 741–751. (doi:10.1038/nrmicro1969)

9. Sinkins SP, Gould F. 2006 Gene drive systems for insect disease vectors. Nat. Rev. Genet. 7, 427–435. (doi:10.1038/nrg1870)

10. Bordenstein SR, O’Hara PF, Werren JH. 2001 *Wolbachia*-induced incompatibility precedes other hybrid incompatibilities in *Nasonia*. Nature 409, 707–710. (doi:10.1038/35055543)

11. Gebiola M et al. 2016 ‘Darwin’s corollary’ and cytoplasmic incompatibility induced by *Cardinium* contribute to speciation in *Encarsia* wasps (Hymenoptera: Aphelinidae). Evolution 70, 2447–2458.

12. Mains JW, Brelsfoard CL, Rose RI, Dobson SL. 2016 Female adult *Aedes albopictus* suppression by *Wolbachia*-infected male mosquitoes. Sci. Rep. 6, 1–7. (doi:10.1038/srep33846)

13. Walker T et al. 2011 The wMel *Wolbachia* strain blocks dengue and invades caged *Aedes aegypti* populations. Nature 476, 450–455. (doi:10.1038/nature10355)

14. Pan X, Zhou G, Wu J, Bian G, Lu P, Raikhel AS, Xi Z. 2012 *Wolbachia* induces reactive oxygen species (ROS)-dependent activation of the Toll pathway to control dengue virus in the mosquito *Aedes aegypti*. Proc. Natl. Acad. Sci. U. S. A. 109, E23–E31. (doi:10.1073/pnas.1116932108)

15. Zug R, Hammerstein P. 2012 Still a host of hosts for *Wolbachia*: Analysis of recent data suggests that 40% of terrestrial arthropod species are infected. PLoS One 7, 7–9. (doi:10.1371/journal.pone.0038544)

16. Sazama EJ, Scot P, Wesner JS. 2019 Insect-symbiont interactions bacterial endosymbionts are common among, but not necessarily within, insect species. Environ. Entomol. 48, 127–133. (doi:10.5063/F1)

17. Takano S et al. 2017 Unique clade of alphaproteobacterial endosymbionts induces complete cytoplasmic incompatibility in the coconut beetle. Proc. Natl. Acad. Sci. 114, 6110–6115. (doi:10.1073/pnas.1618094114)

18. Penz T, Schmitz-Esser S, Kelly SE, Cass BN, Müller A, Woyke T, Malfatti SA, Hunter MS, Horn M. 2012 Comparative genomics suggests an independent origin of cytoplasmic incompatibility in Cardinium hertigii. PLoS Genet. 8. (doi:10.1371/journal.pgen.1003012)

19. Duron O, Cremaschi J, McCoy KD. 2016 The high diversity and global distribution of the intracellular bacterium *Rickettsiella* in the polar seabird tick *Ixodes uriae*. Microb. Ecol. 71, 761–770. (doi:10.1007/s00248-015-0702-8)

20. Curry MM, Paliulis L V, Welch KD, Harwood JD, White JA. 2015 Multiple endosymbiont infections and reproductive manipulations in a linyphiid spider population. Heredity (Edinb). 115, 146–152. (doi:10.1038/hdy.2015.2)

21. Baldo L et al. 2006 Multilocus sequence typing system for the endosymbiont *Wolbachia pipientis*. Appl. Environ. Microbiol. 72, 7098–7110. (doi:10.1128/AEM.00731-06)

22. Kozich JJ, Westcott SL, Baxter NT, Highlander SK, Schloss PD. 2013 Development of a dual-index sequencing strategy and curation pipeline for analyzing amplicon sequence data on the miseq illumina sequencing platform. Appl. Environ. Microbiol. 79, 5112–5120. (doi:10.1128/AEM.01043-13)

23. Caporaso JG et al. 2010 QIIME allows analysis of highthroughput community sequencing data Intensity normalization improves color calling in SOLiD sequencing. Nat. Methods 7, 335–336. (doi:10.1038/nmeth0510-335)

24. Bokulich NA, Subramanian S, Faith JJ, Gevers D, Gordon JI, Knight R, Mills DA, Caporaso JG. 2013 Quality-filtering vastly improves diversity estimates from Illumina amplicon sequencing. Nat. Methods 10, 57–59. (doi:10.1038/nmeth.2276)

25. DeSantis TZ et al. 2006 Greengenes, a chimera-checked 16S rRNA gene database and workbench compatible with ARB. Appl. Environ. Microbiol. 72, 5069–5072. (doi:10.1128/AEM.03006-05)

26. Larsson AJM, Stanley G, Sinha R, Weissman IL, Sandberg R. 2018 Computational correction of index switching in multiplexed sequencing libraries. Nat. Methods 15, 305–307. (doi:10.1038/nmeth.4666)

27. Vanthournout B, Swaegers J. 2011 Spiders do not escape reproductive manipulations by *Wolbachia*. BMC Evol. Biol. 11. (doi:10.1186/1471-2148-11-15)

28. White JA, Kelly SE, Perlman SJ, Hunter MS. 2009 Cytoplasmic incompatibility in the parasitic wasp *Encarsia inaron*: disentangling the roles of *Cardinium* and *Wolbachia* symbionts. Heredity (Edinb). 102, 483–489. (doi:10.1038/hdy.2009.5)

29. Koga R, Tsuchida T, Sakurai M, Fukatsu T. 2007 Selective elimination of aphid endosymbionts: Effects of antibiotic dose and host genotype, and fitness consequences. FEMS Microbiol. Ecol. 60, 229–239. (doi:10.1111/j.1574-6941.2007.00284.x)

30. Williams D. 1982 Extra-binomial variation in logistic linear models. J. R. Stat. Soc. Ser. C (Applied Stat. 31, 144–148.

31. Gotoh T, Noda H, Ito S. 2007 *Cardinium* symbionts cause cytoplasmic incompatibility in spider mites. Heredity (Edinb). 98, 13–20. (doi:10.1038/sj.hdy.6800881)

32. Zhu LY, Zhang KJ, Zhang YK, Ge C, Gotoh T, Hong XY. 2012 *Wolbachia* strengthens *Cardinium*-induced cytoplasmic incompatibility in the spider mite *Tetranychus piercei* McGregor. Curr. Microbiol. 65, 516–523. (doi:10.1007/s00284-012-0190-8)

33. Le Page DP et al. 2017 Prophage WO genes recapitulate and enhance *Wolbachia*-induced cytoplasmic incompatibility. Nature 543, 243–247. (doi:10.1038/nature21391)

34. Beckmann JF, Ronau JA, Hochstrasser M. 2017 A *Wolbachia* deubiquitylating enzyme induces cytoplasmic incompatibility. Nat. Microbiol. 2, 1–7. (doi:10.1038/nmicrobiol.2017.7)

35. Gillespie JJ, Driscoll TP, Verhoeve VI, Rahman MS, Macaluso KR, Azad AF. 2018 A tangled web: Origins of reproductive parasitism. Genome Biol. Evol. 10, 2292–2309. (doi:10.1093/gbe/evy159)

36. Cordaux R, Paces-Fessy M, Raimond M, Michel-Salzat A, Zimmer M, Bouchon D. 2007 Molecular characterization and evolution of arthropod-pathogenic *Rickettsiella* bacteria. Appl. Environ. Microbiol. 73, 5045–5047. (doi:10.1128/AEM.00378-07)

37. Tsuchida T, Koga R, Horikawa M, Tsunoda T, Maoka T, Matsumoto S, Simon J-C, Fukatsu T. 2010 Symbiotic bacterium modifies aphid body color. Science (80-.). 330, 1102–1105. (doi:10.1126/science.1195463.)

38. Kurtti TJ, Palmer AT, Oliver JH. 2002 *Rickettsiella*-like bacteria in *Ixodes woodi* (Acari: Ixodidae). J. Med. Entomol. 39, 534–540. (doi:10.1603/0022-2585-39.3.534)

39. Ochman H, Lawerence JG, Groisman EA. 2000 Lateral gene transfer and the nature of bacterial innovation. Nature 405, 299–304.

40. Ishmael N et al. 2009 Extensive genomic diversity of closely related *Wolbachia* strains. Microbiology 155, 2211–2222. (doi:10.1099/mic.0.027581-0)

41. Goodacre SL, Martin OY, Thomas CFG, Hewitt GM. 2006 *Wolbachia* and other endosymbiont infections in spiders. Mol. Ecol. 15, 517–527. (doi:10.1111/j.1365-294X.2005.02802.x)

42. Ferrari J, West JA, Via S, Godfray HCJ. 2011 Population genetic structure and secondary symbionts in host-associated populations of the pea aphid complex. Evolution (N. Y). 66, 375–390. (doi:10.5061/dryad.8qb00)

43. Russell JA et al. 2013 Uncovering symbiont-driven genetic diversity across North American pea aphids. Mol. Ecol. 22, 2045–2059. (doi:10.1111/mec.12211)

44. Angelella G, Nalam V, Nachappa P, White J, Kaplan I. 2018 Endosymbionts differentially alter exploratory probing behavior of a nonpersistent plant virus vector. Microb. Ecol. 76, 453–458. (doi:10.1007/s00248-017-1133-5)

45. Jaenike J. 2009 Coupled population dynamics of endosymbionts within and between hosts. Oikos 118, 353–362. (doi:10.1111/j.1600-0706.2008.17110.x)

46. Cordaux R, Bouchon D, Grève P. 2011 The impact of endosymbionts on the evolution of host sex-determination mechanisms. Trends Genet. 27, 332–341. (doi:10.1016/j.tig.2011.05.002)

47. Duron O et al. 2017 Evolutionary changes in symbiont community structure in ticks. Mol. Ecol. 26, 2905–2921. (doi:10.1111/mec.14094)

48. Tijsse-Klasen E, Braks M, Scholte EJ, Sprong H. 2011 Parasites of vectors - *Ixodiphagus hookeri* and its *Wolbachia* symbionts in ticks in the Netherlands. Parasites and Vectors 4, 1–7. (doi:10.1186/1756-3305-4-228)

