## Supplementary material for "Endosymbiotic *Rickettsiella* causes cytoplasmic incompatibility in a spider host": Electronic supplemental material figure S1

### Spider gut symbiont detection assay:

#### Methods

We tested whether infected prey could result in false-positive infection diagnoses in *Mermessus fradeorum* using *Wolbachia*-infected *Drosophila melanogaster*. Thirty spiders (10 males, 20 females) from *Wolbachia*-free matrilines were deprived of food for a week before the trial began. Spiders were randomly assigned to one of five treatments: unfed (n=6) or fed a single *D. melanogaster* with DNA extracted one hour (n=6), one day (n=6), three days (n=6), or five days (n=6) after feeding. The six unfed spiders and six positive fruit fly controls were extracted at the same time as the one hour post-feeding treatment. Presence or absence of *Wolbachia* was determined by diagnostic PCR followed by gel electrophoresis of resultant products. Methodology for DNA extraction and PCR were the same as described in the main text.

#### Results

All fruit fly controls yielded strong positive *Wolbachia* bands, but no spiders yielded bands (Figure S1). We conclude that our diagnostic assays are unlikely to yield false positives due to consumption of symbiont-infected prey.

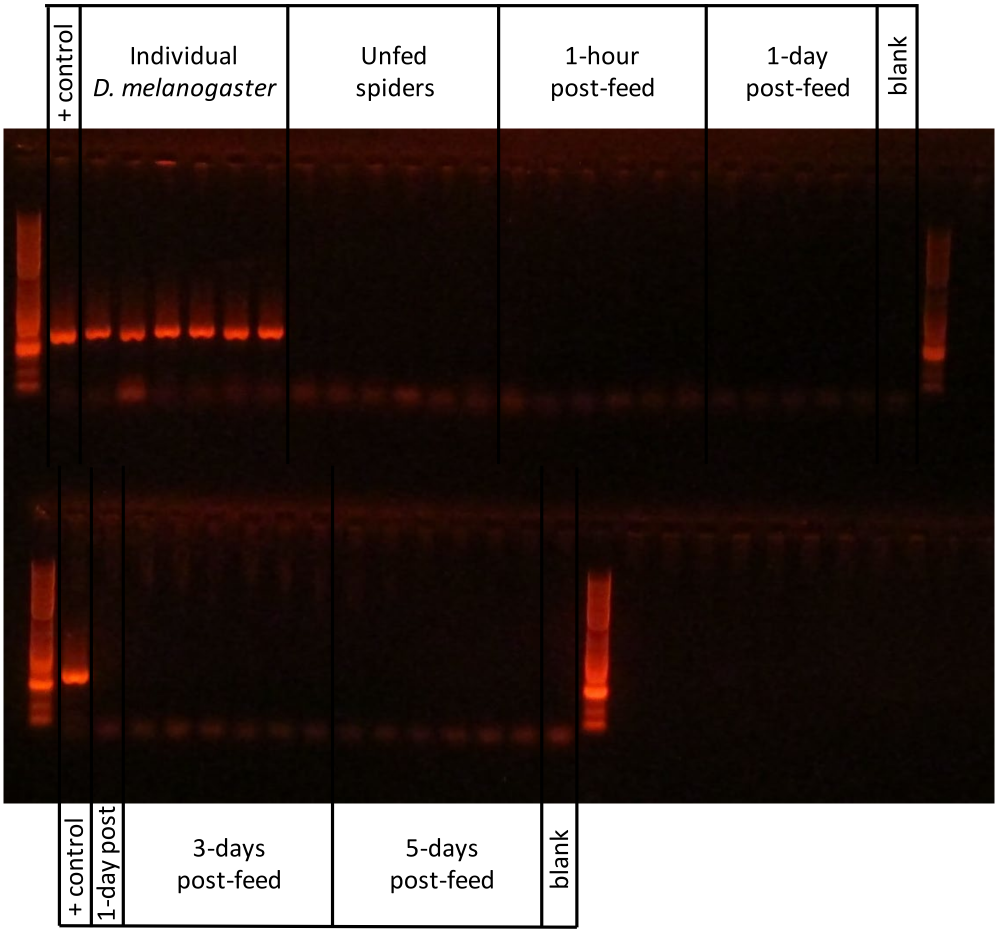


Figure S1. Gel visualization of PCR products from spider gut symbiont detection assay. *Drosophila melanogaster* prey were infected with *Wolbachia* and readily detected by diagnostic *Wolbachia* PCR. However, when the flies were fed to spiders, the *Wolbachia* was undetectable, even in spiders that had been fed a fly only one hour before.
